# A shared brain state for episodic and semantic retrieval

**DOI:** 10.64898/2026.03.25.713662

**Authors:** Matthew B. Bair, Nicole M. Long

## Abstract

It is critical to identify which factors induce specific brain states as these large-scale patterns of coordinated neural activity drive downstream processing and behavior. The retrieval state, a brain state engaged when attempting to retrieve the past, is thought to specifically support episodic memory, remembering experiences within a spatiotemporal context, as opposed to semantic memory, remembering general knowledge. However, we hypothesize that the retrieval state reflects internal attention engaged to access stored episodic and semantic information. To test these alternatives, we recorded scalp electroencephalography while participants made episodic, semantic, or perceptual judgments, and applied an independently validated mnemonic state classifier to measure retrieval state engagement. We found that retrieval state engagement was greater for both episodic and semantic judgments compared to perceptual judgments. These findings suggest that the retrieval state reflects a domain-general internal attention process that supports not just episodic memory, but internally directed cognition.

## Introduction

Imagine being a contestant on the game show Jeopardy! During single, double, and final jeopardy, you must use the answers provided to cue retrieval of some piece of factual information, e.g. “Brenda Milner’s pioneering work helped show that these lobes named for being on the side of the brain are key to memory.” During the “get to know the contestants” break in the middle of single jeopardy, you need to use host Ken Jennings’ question as a cue to retrieve a specific episode from your life. Although in both of these instances the content that you access is different – semantic, factual information vs. episodic, spatiotemporal contextual information^1^ – both likely rely on turning your mind’s eye inward. The retrieval state (or mode^2^) was initially proposed to specifically support episodic, as opposed to semantic, retrieval^3, 4^. However, recent evidence suggests that the retrieval state may reflect a domain-general internal attention process^5–8^ which, if true, would mean that the retrieval state should be engaged during both episodic and semantic retrieval. Identifying the extent to which the retrieval state reflects memory-specific vs. general attention processing is necessary for determining how and when task-relevant and task-irrelevant brain states are recruited to support cognition. The aim of the present study was to investigate the extent to which the retrieval state is recruited during both episodic and semantic retrieval.

The retrieval state is defined as a “tonically maintained cognitive state”^3^ which supports, but is distinct from, retrieval success. Contrasts of both episodic and semantic retrieval tasks^4, 9–13^ and episodic and perceptual tasks^14^ during scalp electroencephalographic (EEG) recordings have revealed a right frontal positivity approximately 600ms following the onset of a task cue and prior to the onset of a stimulus. This right frontal positivity appears to be invariant to the exact episodic retrieval task performed^11^. These findings suggest a spatially-localized, episodic-specific neural correlate of the retrieval state.

However, recent work suggests that the retrieval state extends beyond episodic remembering and is composed of more than a spatially-localized positive voltage deflection. Instead, the retrieval state constitutes a distributed brain state, detectable through multivariate pattern analysis of whole-brain activity. The retrieval state can be distinguished from the encoding state through signals across cortical networks^15^ and across a large range of spectral frequencies^16^. Encoding and retrieval states rely on overlapping, but distinct, hippocampal circuitry such that the two states tradeoff^17, 18^ and engagement in the retrieval state can impair subsequent memory^16, 19^. Furthermore, growing evidence suggests that retrieval is a form of internal attention^5, 6, 8^. For example, the retrieval state tracks the deployment of spatial attention in a non-episodic posner spatial cueing task^7^. Thus, rather than reflect episodic-specific remembering, the retrieval state may map onto the internal axis of attention^5^ whereby internal attention selects among competing internal representations rather than external perceptual input.

Episodic and semantic retrieval rely on overlapping brain networks^20, 21^. Recent evidence has shown that both univariate activation and multivariate patterns are highly overlapping for successful episodic and semantic retrieval^21^. In particular, the default mode network (DMN^22^) is recruited during episodic retrieval^23^, semantic retrieval^24^, and internal mentation more broadly^25^. The DMN is engaged when decisions rely on internally generated, multi-modal information rather than immediate perceptual input, consistent with its role in stimulus-independent cognition^26^. Direct investigation of mnemonic brain states has revealed that the retrieval state recruits a DMN-like microstate, a temporally sustained voltage topography pattern^27^. Together, these findings suggest that the retrieval state may support both episodic and semantic retrieval.

Insofar as engaging the retrieval state impacts downstream processing and behavior^15, 16, 19, 28^, it is necessary to elucidate the types of content and demands that engage the retrieval state. Engaging task-relevant brain states will facilitate behavior whereas engaging task-irrelevant brain states will impair behavior; however, the task-relevance of a given brain state is dependent on the factors that lead to its recruitment. Engaging the retrieval state may impair later memory to the extent that engaging in retrieval prevents encoding of current sensory information^16^. Alternatively, engaging the retrieval state to access stored semantic knowledge to elaborate a memory trace may form the basis of the depth of processing effect^29–31^, facilitating later memory. Thus, optimal mnemonic state engagement – engaging the mnemonic state that is most relevant for current and future goals – is critical for successful behavior across cognition, but optimality will depend on the extent to which these states reflect memory-specific as opposed to broader attentional processes.

Our hypothesis is that both episodic and semantic demands recruit the retrieval state. The alternative hypothesis is that only episodic demands recruit the retrieval state. To test our hypothesis, we conducted a scalp EEG study in which participants studied word stimuli and then completed a three-phase decision task in which they made one of three decisions – old or new (episodic), big or small (semantic), uppercase or lowercase (perceptual) on both old and new words (Figure 1A). We used cross-study multivariate decoding^7, 32, 33^ to assess retrieval state engagement across the three tasks and to link retrieval state engagement with evidence for task and feature information. We expected to find elevated retrieval state evidence during both the episodic and semantic tasks, but not during the perceptual task. All analyses were pre-registered (https://osf.io/jf3ez) unless noted otherwise.

**Figure 1.**
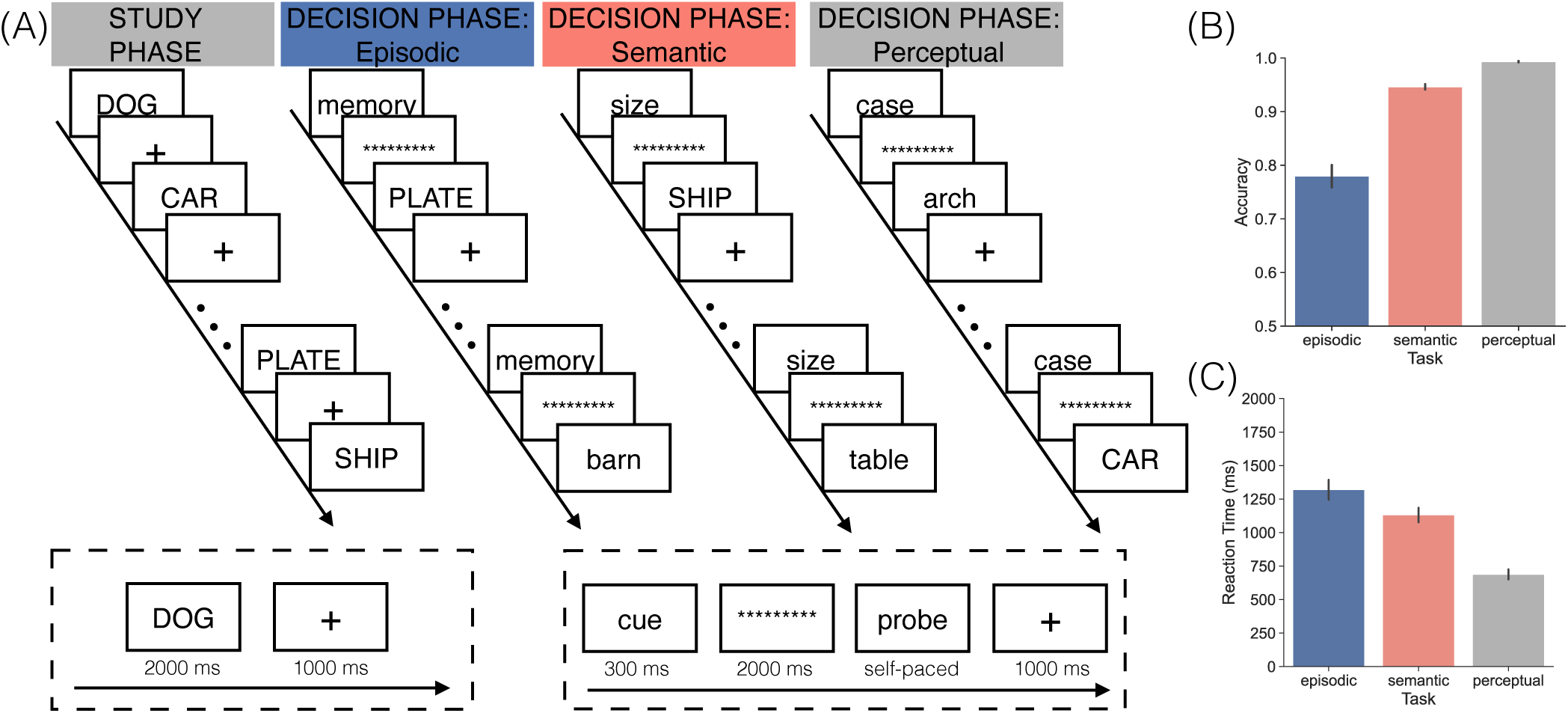
Task design and behavioral results. **(A)** During the study phase, participants viewed eight lists of 24 words for a total of 192 studied words, all presented in uppercase letters. Following the study phase, participants completed the decision phase, which consisted of three blocks of 128 trials, yielding a total of 384 trials across all blocks. Participants performed one task in each of the three blocks. On each trial, a cue appeared on screen indicating the type of judgement that the participants should make about the upcoming probe. In the perceptual task block (“case” cue) participants made an uppercase/lower case judgment. In the semantic task block (“size” cue) participants made a small/big size judgement. In the episodic task block (“memory” cue) participants made an old/new memory judgement. The cue was followed by a row of asterisks, then the probe. Task presentation order and response key mappings were counterbalanced across participants. **(B-C)** Each panel shows the impact of task (episodic, semantic, perceptual) on accuracy **(A)** or reaction time **(B)**. Error bars represent standard error of the mean.

## Results

### Behavioral performance is modulated by task

To examine how task influences behavior, we assessed response accuracy and reaction times (RTs) across the three tasks. We conducted a one-way repeated measures ANOVA (rmANOVA) with task as the factor (episodic, semantic, perceptual) and response accuracy as the dependent variable (Figure 1B). We excluded ambiguous size trials. We find a significant main effect of task on accuracy (*F* _2,56_=79.39, *p<*0.0001, *η_p_*^2^=0.7393). Post-hoc paired *t* -tests between task conditions revealed significantly greater accuracy for the perceptual task (M=0.9933, SD=0.0067) compared to the episodic task (M=0.7796, SD=0.1109; *t* _28_=10.12, *p<*0.0001, *d* =2.720) and the semantic task (M=0.9463, SD=0.0265;*t* _28_=9.048, *p<*0.0001, *d* =2.435). Paired *t* -tests also revealed significantly greater accuracy for the semantic compared to the episodic task (*t* _28_=7.611, *p<*0.0001, *d* =2.068).

We found a similar effect of task on RTs. We conducted a one-way rmANOVA with task as the factor and RT on correct trials as the dependent variable (Figure 1C). We find a significant main effect of task on RTs (*F* _2,56_=85.32, *p<*0.0001, *η_p_*^2^=0.7529). Post-hoc paired *t* -tests between task conditions revealed significantly faster RTs for the perceptual task (M=688.2, SD=198.2) compared to both the episodic (M=1320, SD=391.6; *t* _28_=-10.48, *p<*0.0001, *d* =2.036) and semantic (M=1131, SD=285.4; *t* _28_=-11.69, *p<*0.0001, *d* =1.803) tasks. Paired *t* -tests also revealed significantly faster RTs for the semantic compared to the episodic (*t* _28_=-3.915, *p*=0.0005, *d* =0.5510) task.

In summary, participants performed the perceptual task more accurately and quickly than the semantic task, and the semantic task more accurately and quickly than the episodic task.

### Semantic task demands recruit the retrieval state

Our primary goal was to test the hypothesis that both semantic and episodic demands recruit the retrieval state. To test this hypothesis, we extracted response-locked mnemonic state evidence during the probe interval of the decision phase separately for each task. To the extent that the retrieval state specifically tracks episodic retrieval, we should find positive mnemonic state evidence (greater evidence for the retrieval state) selectively for the episodic task trials. Alternatively, to the extent that the retrieval state tracks content-general internal attention, we should find positive mnemonic state evidence for both the episodic and semantic task trials. We conducted a 3*×*10 rmANOVA with task (episodic, semantic, perceptual) and time interval (100ms time intervals from 500ms preceding the response to 500ms following the response) as factors and response-locked mnemonic state evidence as the dependent variable (Figure 2B). We find a significant main effect of task (*F* _2,56_=14.78, *p<*0.0001, *η_p_*^2^=0.3454) and time interval (*F* _9,252_=17.40, *p<*0.0001, *η_p_*^2^=0.3833). We find a significant interaction between task and time interval (*F* _18,504_=8.181, *p<*0.0001, *η_p_*^2^=0.2261). Thus, both task and time interval modulate mnemonic state engagement during the probe interval.

**Figure 2.**
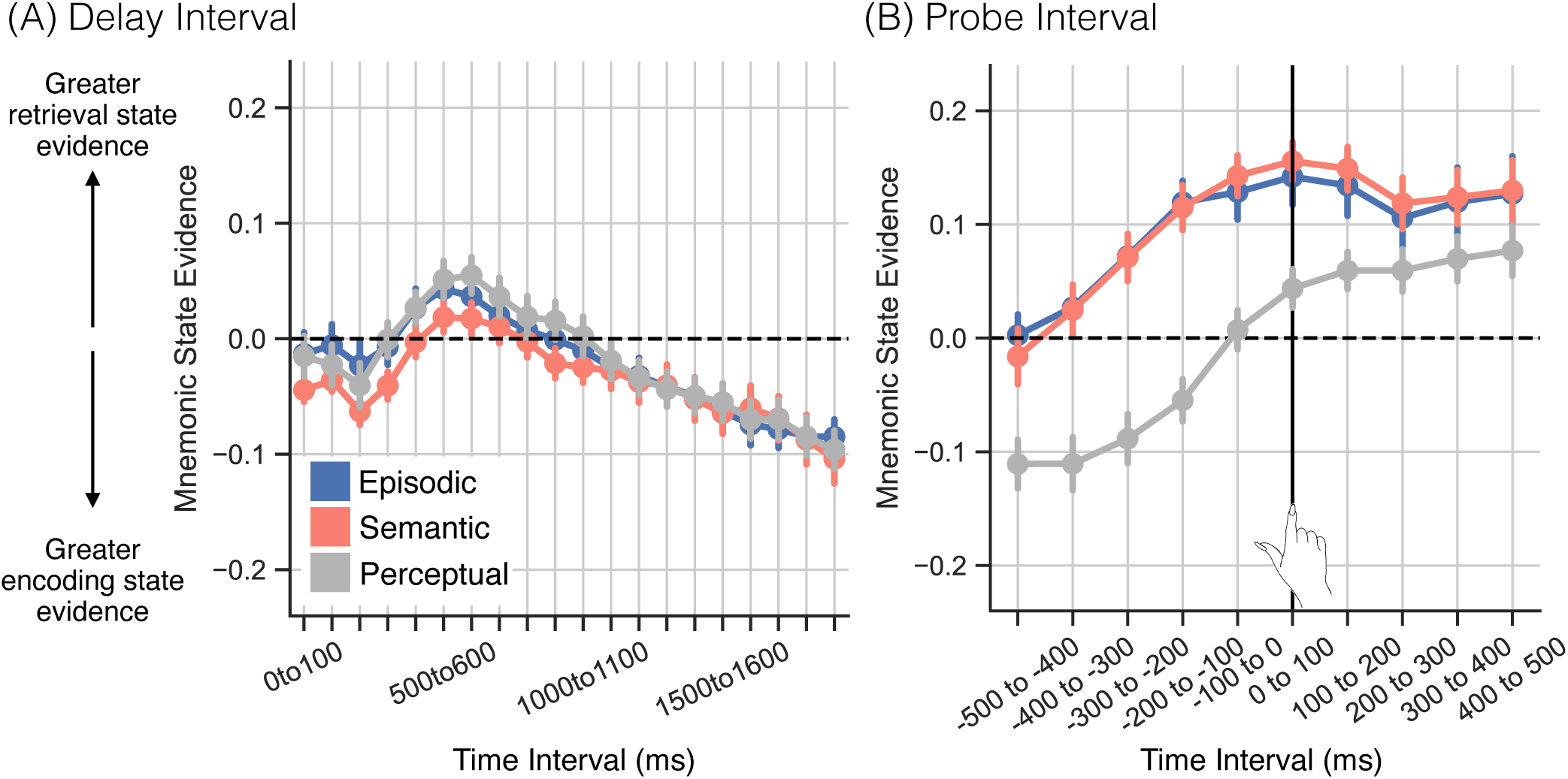
Mnemonic state evidence by task. We applied a cross-study classifier to the decision phase data to measure mnemonic state evidence as a function of task (episodic, blue; perceptual, grey; semantic, red). Positive values indicate greater evidence for the retrieval state; negative values indicate greater evidence for the encoding state. Error bars represent the standard error of the mean. **(A)** We measured mnemonic state evidence across twenty 100 ms time intervals spanning the 2000 ms delay interval. **(B)** We measured response-locked retrieval evidence across ten 100 ms time intervals beginning 500 ms before participant’s response to the probe and ending 500 ms after response to the probe. The vertical line at time 0 to 100 ms indicates the onset of the response.

We conducted pairs of follow-up rmANOVAs to compare each set of tasks. Given our hypothesis that the retrieval state reflects content-general internal attention, we expected to find greater retrieval state engagement for the episodic and semantic tasks compared to perceptual task, and no difference between episodic and semantic tasks. For each set of tasks, we conducted 2*×*10 rmANOVAs with task and time interval as factors (Table 1). For the semantic/perceptual and episodic/perceptual rmANOVAs, we find a main effect of task driven by significantly greater mnemonic state evidence on the semantic (M=0.1014, SD=0.1267) and episodic (M=0.0978, SD=0.1373) compared to the perceptual (M=-0.0048, SD=0.1280) task. For the episodic/semantic rmANOVA, we do not find a significant main effect of task, and Bayes factor analysis revealed that a model without task (H_0_) is preferred to a model with task (H_1_; BF_10_=0.0003, extreme evidence for H_0_).

**Table 1.** Analysis of variance for the effect of task and time interval on mnemonic state evidence across delay and probe intervals.

| Model | Effect | Delay Interval |  |  |  | Probe Interval |  |  |  |
| --- | --- | --- | --- | --- | --- | --- | --- | --- | --- |
| | | df | *F* | *p* | $\eta_p^2$ | df | *F* | *p* | $\eta_p^2$ |
| Semantic x Perceptual | Main effect of task | 1,28 | 1.581 | 0.2190 | 0.0534 | 1,28 | 20.67 | **0.0001** | 0.4247 |
|  | Main effect of time interval | 19,532 | 13.45 | **<0.0001** | 0.3244 | 9,252 | 20.42 | **<0.0001** | 0.4218 |
|  | Interaction of task × time interval | 19,532 | 2.219 | **0.0023** | 0.0734 | 9,252 | 9.505 | **<0.0001** | 0.2534 |
| Episodic x Perceptual | Main effect of task | 1,28 | 1.960 | 0.1720 | 0.0654 | 1,28 | 17.60 | **0.0003** | 0.3859 |
|  | Main effect of time interval | 19,532 | 10.54 | **<0.0001** | 0.2736 | 9,252 | 17.37 | **0.0001** | 0.3829 |
|  | Interaction of task × time interval | 19,532 | 1.816 | **0.0186** | 0.0609 | 9,252 | 11.44 | **<0.0001** | 0.2900 |
| Episodic x Semantic | Main effect of task | 1,28 | 0.0700 | 0.7930 | 0.0025 | 1,28 | 0.0395 | 0.8438 | 0.0014 |
|  | Main effect of time interval | 19,532 | 13.33 | **<0.0001** | 0.3225 | 9,252 | 12.75 | **<0.0001** | 0.3128 |
|  | Interaction of task × time interval | 19,532 | 0.8260 | 0.6760 | 0.0287 | 9,252 | 0.9610 | 0.4729 | 0.0332 |

Bold values indicate $p < 0.05$

Together, these results reveal significantly greater response-locked retrieval state engagement in the semantic and episodic tasks compared to the perceptual task, and no evidence for a difference in retrieval state engagement between the episodic and semantic tasks.

### Retrieval state evidence fluctuates over the delay interval

Early empirical evidence for the retrieval state (e.g.^4^) revealed dissociations between episodic and semantic retrieval tasks during the delay interval between cue and probe. However, recent work from our lab instead suggests that the retrieval state is selectively recruited following probe onset, rather than in response to the instruction cue^19^. Insofar as the retrieval state reflects content-general internal attention, it should be recruited for all three tasks during the delay interval to support the selection of task sets^5^.

We assessed delay-interval mnemonic state engagement across the three tasks. We did not expect to find an effect of or interaction with task. We anticipated that mnemonic state evidence might change across the delay interval. Specifically, we expected that early in the delay interval there may be a task-general increase in mnemonic state evidence (reflecting retrieval state engagement) followed by a task-general decrease in mnemonic state evidence (reflecting encoding state engagement). Retrieval state engagement would putatively reflect selection of the relevant task set, whereas encoding state engagement would putatively reflect a preparatory response in anticipation of the upcoming probe. We conducted a 3*×*20 rmANOVA with task (episodic, semantic, perceptual) and time interval (100ms time intervals spanning the 2000ms delay interval) as factors, and mnemonic state evidence as the dependent variable (Figure 2A). We do not find a significant main effect of task (*F* _2,56_=1.125, *p*=0.3319, *η_p_*^2^=0.0386). We find a significant main effect of time interval (*F* _19,532_=13.37, *p<*0.0001, *η_p_*^2^=0.3231) and a significant interaction between task and time interval (*F* _38,1064_=1.587, *p*=0.014, *η_p_*^2^=0.0536).

As a result of the task by instruction interaction, we conducted pairs of follow-up 2*×*20 rmANOVAs to compare each set of tasks (Table 1). For the semantic/perceptual and episodic/perceptual rmANOVAs, we find a significant interaction between task and time interval, driven by an overall shift in mnemonic state evidence between the semantic/episodic tasks compared to the perceptual task within the first 1000ms of the delay interval. Mnemonic state evidence is more positive for episodic and semantic tasks relative to the perceptual task. However, for both comparisons, Bayes factor analysis revealed that a model without task (H_0_) is preferred to a model with task (H_1_; semantic/perceptual: BF_10_=0.1378, moderate evidence for H_0_; episodic/perceptual: BF_10_=0.0107, very strong evidence for *H*_0_). For the episodic/semantic rmANOVA, we do not find a significant main effect of task or a significant interaction between task and time interval. Bayes factor analysis revealed that a model without task (H_0_) is preferred to a model with task (H_1_: BF_10_*<*0.0001, extreme evidence for H_0_).

Given the relatively modest task effects during the delay interval, we averaged mnemonic state evidence across task and conducted post-hoc one sample comparisons at each time point in the delay interval (Table 2). We find that mnemonic state engagement changes across the delay interval with significant increases from 500-700ms and significant decreases from 200-300ms and from 1200-2000ms following delay interval onset. Our interpretation is that regardless of task, the delay interval begins with continued external processing of the (now absent) cue stimulus, followed by retrieval state engagement to access or prepare the appropriate task set, with a gradual shift to the encoding state in anticipation of the upcoming probe stimulus.

**Table 2.** One sample comparisons of mnemonic state evidence during the delay interval.

| **Time Interval (ms)** | **$t_{28}$** | **$p$** | **$\eta_p^2$** |
| --- | --- | --- | --- |
| 0 to 100 | -2.2242 | 0.0344 | 0.4203 |
| 100 to 200 | -1.9467 | 0.0617 | 0.3679 |
| **200 to 300** | **-3.2462** | **0.0030** | **0.6135** |
| 300 to 400 | -1.678 | 0.1044 | 0.3172 |
| 400 to 500 | 1.552 | 0.1318 | 0.2934 |
| **500 to 600** | **3.359** | **0.0023** | **0.6348** |
| **600 to 700** | **3.329** | **0.0025** | **0.6291** |
| 700 to 800 | 1.8805 | 0.0705 | 0.3554 |
| 800 to 900 | 0.6157 | 0.5431 | 0.1164 |
| 900 to 1000 | -0.1812 | 0.8575 | 0.0342 |
| 1000 to 1100 | -0.8856 | 0.3834 | 0.1674 |
| 1100 to 1200 | -1.718 | 0.0968 | 0.3247 |
| **1200 to 1300** | **-2.464** | **0.0201** | **0.4657** |
| **1300 to 1400** | **-3.002** | **0.0056** | **0.5674** |
| **1400 to 1500** | **-3.458** | **0.0018** | **0.6534** |
| **1500 to 1600** | **-4.118** | **0.0003** | **0.7782** |
| **1600 to 1700** | **-4.268** | **0.0002** | **0.8067** |
| **1700 to 1800** | **-4.581** | **0.0001** | **0.8657** |
| **1800 to 1900** | **-5.293** | **<0.0001** | **1.000** |
| **1900 to 2000** | **-6.035** | **<0.0001** | **1.141** |

Bold values indicate tests that survive FDR correction.

### Task information is decodable during both the delay and probe intervals

To perform the three decision phase tasks, participants should leverage task information such as response mappings, cue specifications and other high-level or abstracted processes^34–37^. Thus it should be possible via multivariate activity patterns to decode the task as participants prepare (delay interval) and/or perform (probe interval) said task^38–40^. Such task evidence may be linked to engagement of the retrieval state insofar as this state reflects internally directed attention to abstract goals and task sets.

To first determine whether task is decodable, we performed leave-one-participant-out cross-validated classification analyses on spectral power averaged across either the 2000ms delay interval (Figure 3A) or the 500ms preceding the response during probe interval (Figure 3B). We used paired-sample *t* -tests to compare classification accuracy across participants to chance decoding accuracy (as determined by permutation procedure, see Methods). We find significantly above chance decoding during both the delay interval (M=36.24%, SD=1.600%; *t* _28_=9.502, *p<*0.0001, *d* =2.531) and probe interval (M=44.60%, SD=1.950%; *t* _28_=30.01, *p<*0.0001, *d* =8.179).

**Figure 3.**
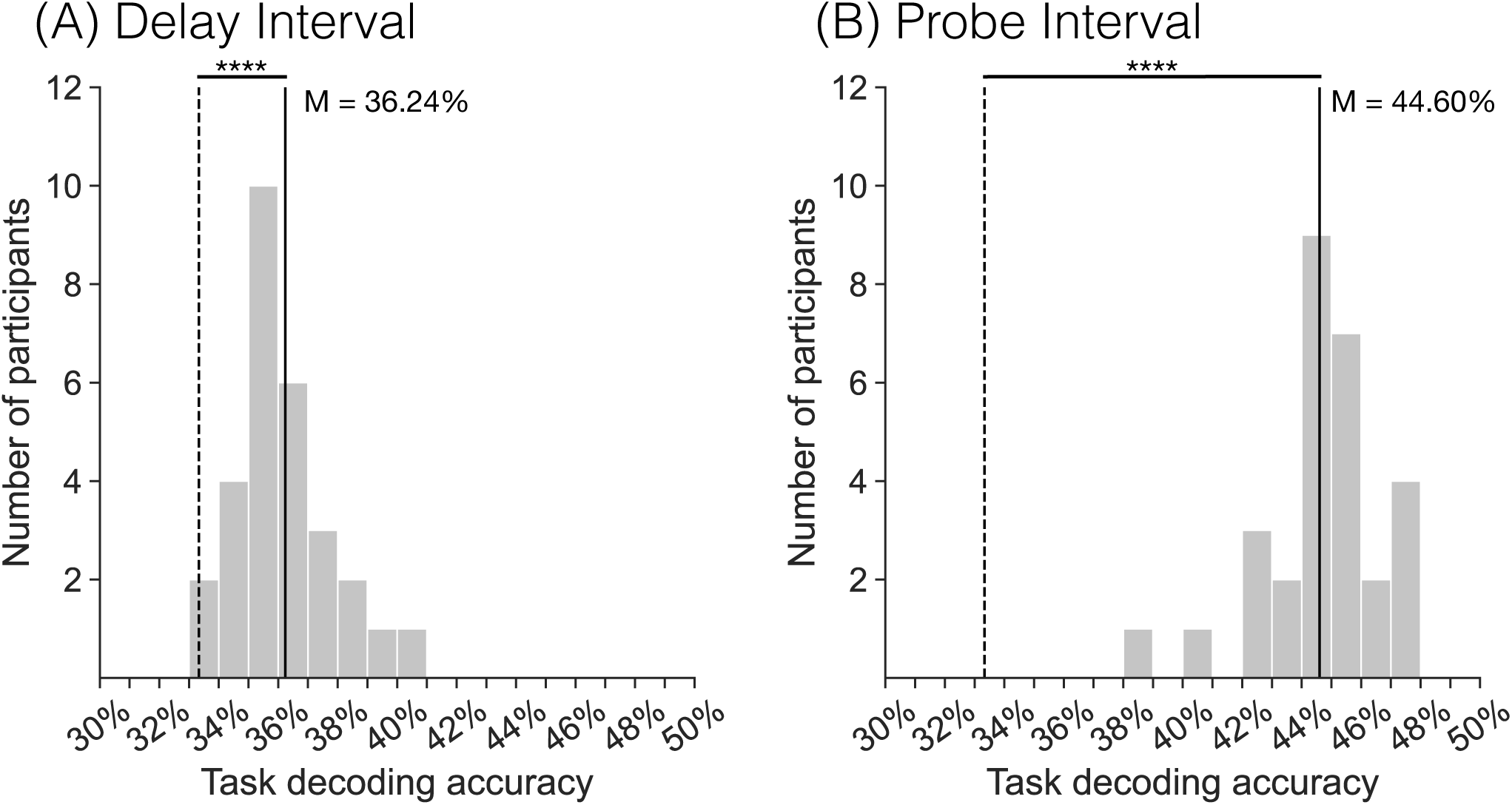
Cross-participant task decoding accuracy. Each panel shows the distribution of cross-participant task-decoding accuracy values for either the 2000 ms delay interval **(A)** or the 500 ms interval preceding the participants’ response during the probe interval **(B)**. We used leave-one-participant-out cross-validated classification to decode the three decision phase tasks (episodic, perceptual, semantic). Solid black vertical lines indicate mean classification accuracy, as determined by permutation procedure. Dashed black vertical lines indicate chance classification accuracy. ****p*<*0.0001.

To the extent that common task-related information is present during both the delay and probe intervals, we should find that the task classifier “generalizes” such that cross-interval decoding is significantly above chance. To test this possibility, we conducted two leave-one-participant-out cross-validated classification analyses, one in which the classifier was trained on delay interval data and tested on probe interval data and one in which the classifier was trained on probe interval data and tested on delay interval data. We used paired-sample *t* -tests to compare classification accuracy across participants to chance decoding accuracy as determined by permutation procedure. We find significantly above chance decoding when the classifier is trained on delay interval data and tested on the probe interval data (M=36.32%, SD=1.160%; *t* _28_=13.56, *p<*0.0001, *d* =3.649), and when the classifier is trained on the probe interval data and tested on the delay interval data (M=38.31%, SD=1.530%; *t* _28_=17.23, *p<*0.0001, *d* =4.617).

In summary, task identity was successfully decoded during both the delay interval and probe intervals, and the above chance cross-interval decoding suggest that these intervals may share common task information.

### Feature information unfolds over time and in response to task relevance

In addition to task-related information, participants should leverage feature information during the probe interval. Every probe in our experiment had three features: old or new, small or big, and uppercase or lowercase. One feature was task-relevant and the remaining two were task-irrelevant. For example, during the episodic task, old/new features were task-relevant and small/big, uppercase/lowercase features were task-irrelevant. In line with our framework that both episodic and semantic tasks recruit internal attention, the episodic (old/new) and semantic (small/big) features should be considered ‘internal’ features. In contrast, the perceptual features (uppercase/lowercase) should be considered ‘external’ features – features readily available to sensory perception. We expected that task relevance would differentially impact the decoding of putative external vs. internal features. Prior work has demonstrated that stimulus features are decodable, especially and sometimes exclusively when said features are task-relevant (e.g.^40–44^). We predicted that internal features would only be decodable when task-relevant, whereas external features would be decodable regardless of task relevance. Furthermore, we view time as especially critical to the differentiation between external and internal features: external features should be decodable soon after probe onset – that is, early during the probe interval – whereas internal feature decoding should be delayed, with internal features decodable only after the probe stimulus has been perceived.

We conducted six cross-validated within-participant pattern classification analyses on spectral power averaged across the 500ms preceding the response during the probe interval. We used paired-sample *t* -tests to compare classification accuracy across participants on task-relevant and task-irrelevant classifiers to chance decoding accuracy and to directly compare classification accuracy across task-relevant and task-irrelevant trials (Figure 4; Table 3). Both episodic and perceptual features were decodable during both task-relevant and task-irrelevant trials (Figure 4A, C). In contrast, semantic features were only decodable during task-relevant trials (Figure 4B). For all three sets of features, we found significantly higher decoding accuracy for task-relevant vs. task-irrelevant trials.

**Figure 4.**
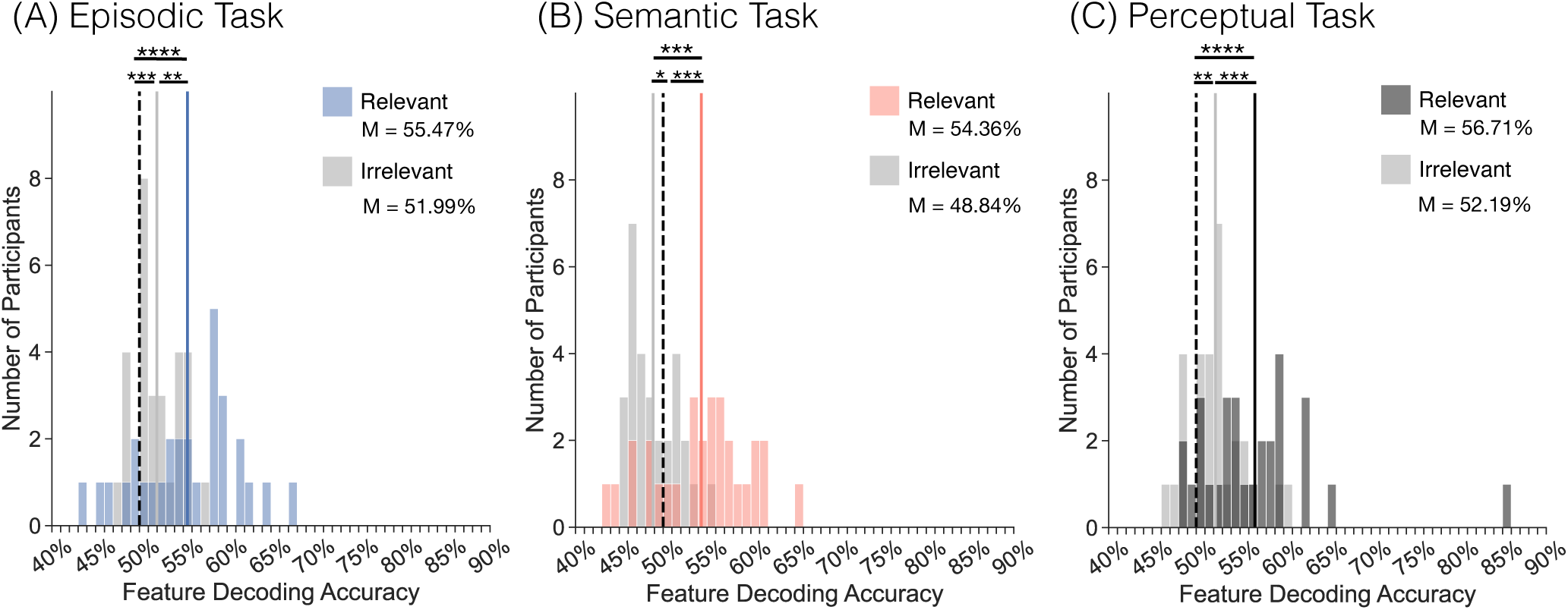
Within-participant feature decoding accuracy. Each panel shows the distribution of within-participant, feature decoding accuracy values for both task-relevant (**A** episodic, blue; **B** semantic, red; **C** perceptual, dark grey, respectively) and task-irrelevant decoders (light grey). We ran six within-participant classifiers which were trained to distinguish task features (old/new, small/big, or uppercase/lowercase) based on response-locked signals in the 500 ms interval preceding participants’ response to the probe. Solid vertical lines indicate mean classification accuracy. Dashed vertical lines indicate chance classification accuracy as determined by permutation procedure. **(A-C)** For all three tasks, decoding accuracy was significantly greater for the task-relevant vs. task-irrelevant decoder. *p*<*0.05, **p*<*0.01, ***p*<*0.001, ****p*<*0.0001.

**Table 3.**
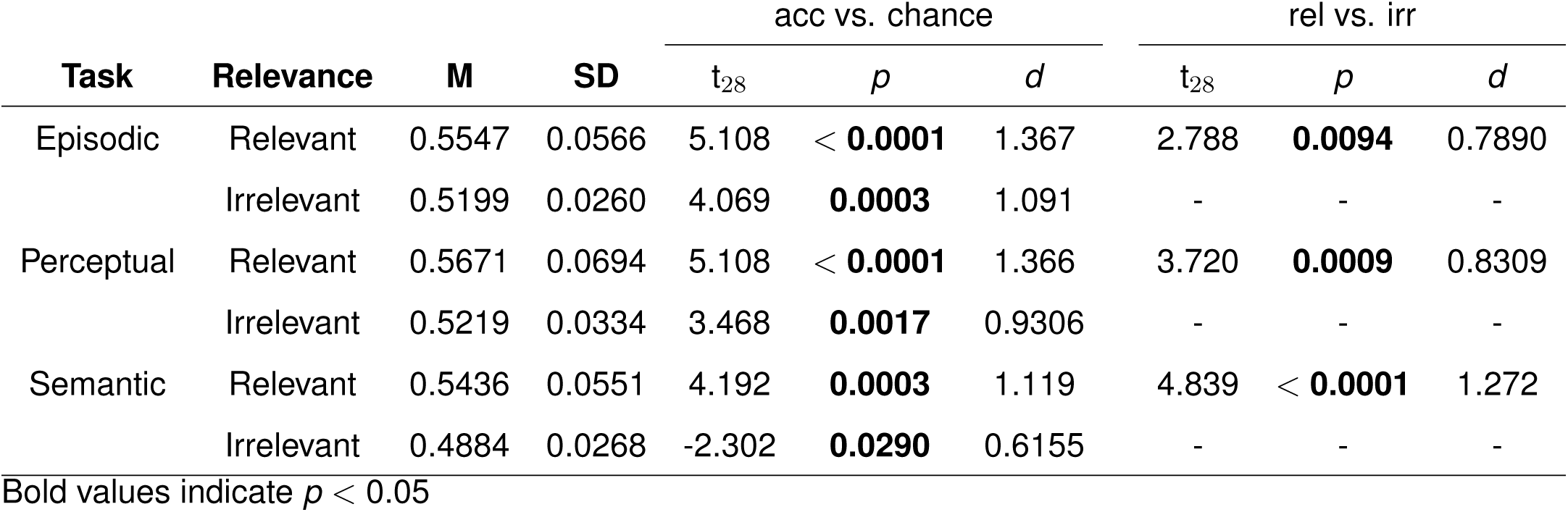
Feature decoding accuracy.

| Task | Relevance | M | SD | acc vs. chance |  |  | rel vs. irr |  |  |
| --- | --- | --- | --- | --- | --- | --- | --- | --- | --- |
| | | | | $t_{28}$ | $p$ | $d$ | $t_{28}$ | $p$ | $d$ |
| Episodic | Relevant | 0.5547 | 0.0566 | 5.108 | < **0.0001** | 1.367 | 2.788 | **0.0094** | 0.7890 |
|  | Irrelevant | 0.5199 | 0.0260 | 4.069 | **0.0003** | 1.091 | - | - | - |
| Perceptual | Relevant | 0.5671 | 0.0694 | 5.108 | < **0.0001** | 1.366 | 3.720 | **0.0009** | 0.8309 |
|  | Irrelevant | 0.5219 | 0.0334 | 3.468 | **0.0017** | 0.9306 | - | - | - |
| Semantic | Relevant | 0.5436 | 0.0551 | 4.192 | **0.0003** | 1.119 | 4.839 | < **0.0001** | 1.272 |
|  | Irrelevant | 0.4884 | 0.0268 | -2.302 | **0.0290** | 0.6155 | - | - | - |

Bold values indicate $p < 0.05$

Having shown that probe features are decodable, we next sought to determine when during the probe interval each feature type (e.g. external, internal) was maximally decodable. We expected that the putative external feature of probe word case would reach the highest decoding accuracy earlier than the two putative internal features of episodic status and size. That is, before you can decide whether a “SHIP” is small or big, you must first process low-level visual features and identify the word that is being presented. Using probe interval, stimulus-locked data, we assessed the time interval at which decoding accuracy was highest for each feature, separately for task-relevant and task-irrelevant trials (Figure 5). We excluded task-irrelevant semantic features given that they were not decodable in the response-locked analysis above.

**Figure 5.**
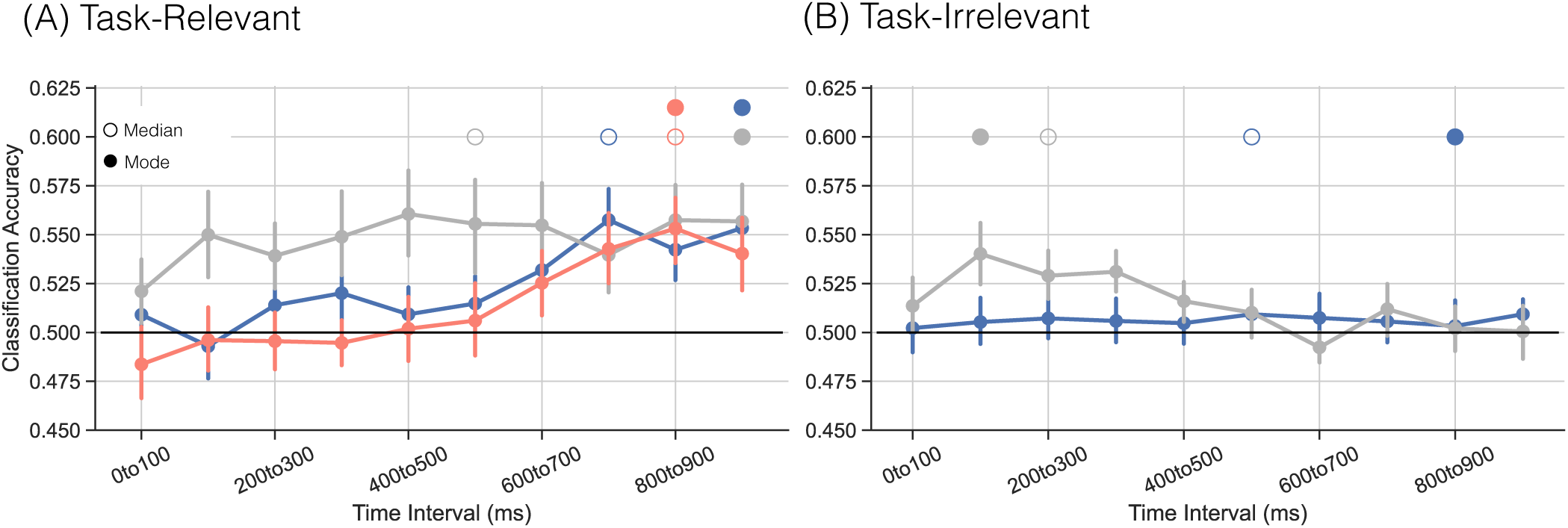
Feature decoding accuracy across the probe interval. Panels show classification accuracy for either task-relevant **(A)** or task-irrelevant **(B)** feature decoding classifiers applied to probe-locked data during the probe interval. We extracted peak decoding time for each participant and calculated median (unfilled circles) and mode (filled circles) peak times across participants. Horizontal lines indicate chance-level classification accuracy. Error bars represent the standard error of the mean. As task-irrelevant semantic feature decoding accuracy did not exceed chance, it is not included in **(B)**.

We extracted the median peak feature decoding time interval for each task-relevant feature. We used Wilcoxon rank-sum tests to compare the peak time interval across features. The peak decoding interval for perceptual features (*Mdn*=500-600ms) occurred significantly earlier than for semantic features (*Mdn*=800-900ms; *z*_28_=-2.115, *p*=0.0344, r=0.2810) and numerically earlier than for episodic features (*Mdn*=700-800ms; *z*_28_=-1.788, p=0.0737, r=0.2380). We did not find a significant difference in peak time between episodic and semantic features (*z*_28_=-0.5832, *p*=0.5598, r=0.0781).

We had anticipated that external features would be decodable regardless of task relevance and indeed, we found above chance decoding of perceptual features during the perceptual task trials as well as during the episodic and semantic task trials. However, we speculated that the temporal dynamics of this feature information may vary as a function of task-relevance, with task-irrelevant perceptual features being maximally decodable even earlier in the probe interval. We directly compared task-relevant and task-irrelevant trials and found that peak decoding time for task-irrelevant perceptual features (*Mdn*=200-300ms) occurred significantly earlier than for task-relevant perceptual features (*z*_28_=-2.683, *p*=0.0073, r=0.356).

Together these findings present a nuanced picture of how feature information is coded. Consistent with prior work, task-relevant features are more strongly decodable than task-irrelevant features. However, the time course of feature information varies depending on both the type and task-relevance of the specific feature. Internal features, or information not overtly present in the sensory properties of the stimulus itself, are maximally decodable nearly 1000ms after the onset of the probe stimulus. External features, here the case of the presented word, are maximally decodable several hundred milliseconds earlier than internal features when task-relevant and even earlier when task-irrelevant. We speculate that external features are initially automatically activated regardless of relevance – likely in very early sensory/perceptual regions – but that external feature information diminishes if not task-relevant. In contrast, internal features require more time to become activated and only do so after a cascade of low-level into higher-level processing.

### Linking mnemonic states with task and feature information

Our final goal was to directly link mnemonic state evidence with task and feature information. To the extent that mnemonic states track internal and external attention, mnemonic state evidence may scale with either or both task and feature information. Insofar as participants select a task set during the delay interval, and such selection recruits internal attention^5^, we should find a positive association between trial-level task evidence and mnemonic state evidence across all three tasks. Alternatively, mnemonic states may be linked with stimulus features rather than abstract task sets, in which case we may fail to find an association between task and mnemonic state evidence. Finally, there may be a negative association whereby decreased task evidence is associated with increased mnemonic state evidence, reflecting greater engagement of the retrieval state in the attempt to access the stored task set information.

To reduce the number of possible comparisons, we extracted a single value for task evidence and mnemonic state evidence for the delay interval (by averaging power across the 2000ms interval and testing each classifier on this averaged data). We performed a Pearson correlation on the trial-level evidence values separately for the three tasks and averaged z-Rho values across the tasks. We did not find a significant correlation between retrieval state and task evidence during the delay interval (M=-0.0201, SD=0.1786; *t* _28_=-1.465, *p*=0.1540, *d* =0.2769).

The lack of a correlation between delay-interval task and retrieval state evidence is not surprising given that we observed relatively low (albeit above chance) task decoding and retrieval state modulation during this interval. Given the robust effects during the probe interval, we conducted an exploratory analysis to test the association between response-locked retrieval state and task evidence. We used the same methods described above and extracted evidence values from a single response-locked time window from the probe interval (−500 to 0ms). We found a significant positive correlation (M=0.0576, SD=0.3424; *t* _28_=6.155, *p<*0.0001, *d* =1.163) whereby greater retrieval state evidence is associated with greater task evidence.

In addition to a probe-interval association between task and retrieval state evidence, there may be a link between feature and retrieval state evidence. We have previously found that retrieval state evidence is positively correlated with a maintained probe’s spatial location^7^ and could expect a similar association for task-relevant, internal features. To reduce the number of possible comparisons, we extracted a single value for feature and mnemonic state evidence during the probe interval (by averaging power across the 500ms pre-response interval and testing each classifier on this averaged data). We performed a Pearson correlation on trial-level evidence values for the four features (old, new, big, small). After Fisher-Z transforming the rho values, we averaged across all features. We did not find a significant correlation between internal features and retrieval evidence (M=-0.0045, SD=0.1603; *t* _28_=-0.3525, *p*=0.7271, *d* =0.0666).

Finally, to the extent that participants focus on external features, we may find a negative association between external feature evidence and mnemonic state evidence during the probe interval. Due to the design of our classifier, negative mnemonic state evidence values reflect evidence for the encoding state. Therefore, we predicted that greater external feature evidence (uppercase/lowercase) would be associated with greater evidence for encoding. To reduce the number of possible comparisons, we extracted a single value for feature and mnemonic state evidence during the probe interval. We performed a Pearson correlation on trial-level evidence values for the two features (uppercase, lowercase). After Fisher-Z transforming the rho values, we averaged across all features. We did not find a significant correlation between external features and retrieval evidence (M=-0.0038, SD=0.1493; *t* _28_=-0.2231, *p*=0.8251, *d* =0.0422).

In summary, we found a significant positive association between trial-level task and mnemonic state evidence during the probe interval and no significant associations between internal or external feature information and mnemonic state evidence.

## Discussion

The goal of this study was to assess the degree to which episodic and semantic task demands recruit the retrieval state. Using scalp EEG and multivariate decoding, we measured memory brain state engagement while participants performed perceptual, episodic, or semantic tasks. Consistent with our hypothesis, we found that both the episodic and semantic tasks strongly recruited the retrieval state whereas the perceptual task did not. Retrieval state recruitment was positively linked with evidence for the task being performed. We found that episodic and semantic stimulus features were maximally dissociable later in time compared to perceptual stimulus features. Together, these findings suggest that the retrieval state reflects domain-general internal attention that flexibly supports goal-directed processing across both episodic and semantic retrieval.

We find that the retrieval state is engaged when participants make both episodic (recognition) and semantic (size) judgments relative to perceptual (case) judgments during presentation of a probe stimulus. We find limited evidence for task-based mnemonic state dissociations during the delay interval. Instead, retrieval state evidence positively correlates with task evidence during the probe interval. These findings run counter to early work demonstrating a spatially-localized episodic retrieval mode characterized by a voltage positivity over right frontal cortex that is invoked in preparation for upcoming episodic retrieval^4, 9–14^. We speculate, especially given task decoding effects, that prior work differentiated task set components. At least one participant indicated that they imagined a shoebox during the semantic, size judgment. Thus, previous episodic vs. semantic dissociations may reflect differences in visualization related to task goals, rather than memory-related processing dissociations. Our findings provide strong support for our hypothesis that the retrieval state reflects content-general internal attention rather than episodic-specific remembering. Internal attention should be engaged when any stored information is accessed, whether a distinct event from a specific time and place or general factual knowledge. Furthermore, internal attention requires a target upon which one focuses the mind’s eye^5, 19^ consistent with our finding of robust mnemonic state dissociations during the probe interval when a putative target for attention is present.

Corroborating our interpretation that episodic and semantic retrieval recruit internal attention, we find that episodic and semantic stimulus feature decoding is delayed relative to perceptual features. Prior work has shown that stimulus features are selectively decodable when task-relevant^40–44^. We provide an important advance to this work by demonstrating that the time course of feature decoding varies as a function of multiple properties. First, features that are external or directly perceivable – e.g. the case of the words presented – are decodable earlier than putative internal features, or those that are not directly perceivable – e.g. whether the word represents an object that is bigger or smaller than a shoebox. These findings are consistent with a framework in which stimulus processing proceeds from low-level visual/perceptual input to high-level recognition, categorization, and extraction of abstract information^45, 46^. Second, with the high temporal resolution of scalp EEG, we were able to detect fine-scale temporal dynamics of feature information and reveal that external features are decodable even when task-irrelevant, but only very early in the probe interval. This finding provides new insights into how feature information is represented, with implications for how potentially task-irrelevant, but salient distracting features may impact attention. Finally, we find that episodic features are decodable when task-irrelevant, meaning that the brain differentiates repeated from novel items regardless of goals. This finding is consistent with prior work suggesting that the retrieval state may be automatically recruited on the basis of bottom-up stimulus factors such as repetition and familiarity^8, 32, 47^. Together, these results show that stimulus feature representations unfold over time: external features are represented early and are maintained only when task-relevant, whereas internal features emerge later, reflecting delayed processing of high-level information.

That the retrieval state is recruited for both episodic and semantic tasks is in line with growing evidence that suggests a strong degree of overlap between episodic and semantic retrieval. Both forms of retrieval are known to recruit the default mode network (DMN;^20, 22, 24^), a network that generally supports internal mentation^25^. Recent work has shown that both univariate activity and multivariate activity patterns are largely similar across episodic and semantic retrieval^21^. Here we show that a shared multivariate brain state supports both forms of retrieval.

Linking neural signals to cognition depends on the underlying brain states^48^ as do efforts to use stimulation to alter neural signals and behavior^49^. Thus, to understand and establish optimal brain state engagement, it is first necessary to identify the factors that induce brain states. Engaging the retrieval state during stimulus presentation can interfere with encoding^16, 19^. However, given growing evidence that the retrieval state reflects internal attention rather than episodic memory retrieval specifically^6–8^, engaging in the retrieval state may not always be detrimental for later memory. Although external attention should support memory for perceptual information in the environment, internal attention may support the formation of semantically elaborated memories^29, 30^. The factors that induce mnemonic and attentional brain states serve as boundary conditions that establish what these states reflect, in turn informing when and how brain states should be optimally engaged.

In conclusion, we show that the retrieval state is engaged during both episodic and semantic judgments, indicating that it is not specific to episodic memory retrieval. These findings support the interpretation that the retrieval state reflects a domain-general internal attention state. Given the role of internal processing across many domains including perception, attention, memory, and decision making, this brain state is expected to have broad impact on behavior across cognition.

## Methods

### Participants

Thirty-four adult (18 female, age range=18-45, mean age=23 years) fluent English speakers from the University of Virginia (UVA) community participated. Our sample size was determined a priori based on behavioral pilot data (N=11) described in the pre-registration report of this study (https://osf.io/jf3ez). Participants were recruited from flyers posted around the UVA campus and surrounding areas. All participants had normal or corrected-to-normal vision. Informed consent was obtained in accordance with the UVA Institutional Review Boards for Social and Behavioral Research and participants were compensated for their participation. Five participants were excluded from the final dataset: One participant who did not complete the experiment, two participants who failed to comply with instructions, and two participants who performed poorly on the perceptual or semantic tasks (response accuracy *<* 2.5*SDs of the mean of the full dataset). We report data for the remaining 29 participants. The raw, de-identified data and the associated experimental and analysis codes used in this study will be made available via the Long Term Memory laboratory website (https://longtermmemorylab.com) upon publication.

### Materials

Stimuli consisted of 384 words reflecting common objects found in the Oliva object category database (e.g. bench, fence; http://olivalab.mit.edu/MM/objectCategories.html). We selected stimuli that enable small/big responses to a size judgment (see below). We eliminated objects delineated by two word terms (e.g. “bottle opener”) and ambiguous objects (e.g. “bat” which could refer to the sporting equipment or the animal).

### Experimental Design

#### Study Phase

Participants studied lists of words. Participants then completed a three-block “decision phase” in which they performed one of three tasks and made one of two judgments in each task (Figure 1A). We blocked trials by task to maximize stay trials to facilitate detection of mnemonic state engagement^12^.

#### Study Phase

In each of eight runs, participants viewed a list containing 24 words, yielding a total of 192 trials. Each study trial consisted of a single word presented in uppercase on a white background. Each word was presented for 2000 ms and was followed by a 1000 ms inter-stimulus interval (ISI). Participants made no behavioral responses during the study phase.

#### Decision Phase

Following the final study list, participants completed the decision phase. The decision phase consisted of three blocks. Each block consisted of 128 trials in which a single word was presented on screen, yielding a total of 384 trials across the decision phase. Each word fit into one of eight possible conditions based on probe (target, lure), case (uppercase, lowercase), and size of the object represented by the word (small, big). Within each block, half of the words were targets (presented during the study phase) and half of the words were lures (not presented during the study phase). Half of the words were be presented in uppercase and half were presented in lowercase. Size was not strictly be balanced, see more below.

Participants performed one of three tasks in each of the blocks. In the episodic task, participants made an old/new judgment and responded as to whether the presented word had or had not been presented during the study phase. In the perceptual task, participants made a case judgment and responded as to whether the presented word was in uppercase or lowercase. In the semantic task, participants made a size judgment and responded as to whether the presented word represents a stimulus that can (small) or cannot (big) fit inside a shoebox. Task order was randomly assigned for each participant. Prior to the start of each block, participants were cued as to which task to perform and were reminded of response mappings. Trials in each block proceeded as follows. The trial began with a cue (episodic, “memory;” semantic, “size;” perceptual, “case”) presented for 300 ms. The cue was followed by a 2000 ms delay interval during which a row of asterisks was presented. Following the delay interval, the probe word was presented until participants responded, thus responses were self-paced. With this design, we followed the prior literature^11–14, 50^ with one exception. In prior work, although participants’ responses were self-paced, the probe itself was presented for a limited amount of time, 300 ms. We modified this aspect of the design based on our prior work and predictions for retrieval state engagement. Specifically, we have found elevated retrieval state engagement during the response interval when the to-be-responded to probe is no longer visually presented^7^. We anticipated that such an effect would be found for all decision types (episodic, semantic, perceptual) insofar as participants would have to access the internal, mental representation of the just-presented-probe in order to make their decision. Thus, we modified the design so that the probe remained visually presented throughout the response interval. Participants responded with their left and right hand via the d and k keys. Response mappings were counterbalanced across participants such that all possible combinations of responses (eight in total) were experienced across all participants. Participants were encouraged to respond as quickly and accurately as possible.

#### Size Judgments

We obtained size ratings for the 384 words from the 28 participants who were included in the behavioral pilot version of this experiment. To assign a label of either small or big, we required 75% agreement across the pilot participants who evaluated a particular stimulus (not all participants made size judgments about all stimuli). With this threshold, we obtained 186 small items, 127 big items, and 71 ambiguous items. Ambiguous items were included in order to reach to the total of 384 items, but we excluded them from all analyses.

### EEG Data Acquisition and Preprocessing

All acquisition and preprocessing methods are based on our previous work^28^; for clarity we use the same text as previously reported. Electroencephalography (EEG) recordings were collected using a BrainVision system and an ActiCap equipped with 64 Ag/AgCl active electrodes positioned according to the extended 10–20 system. All electrodes were digitized at a sampling rate of 1000 Hz and were referenced to electrode FCz. Offline, electrodes were later converted to an average reference. Impedances of all electrodes were kept below 50 kΩ. Electrodes that demonstrated high impedance or poor contact with the scalp were excluded from the average reference calculations; however, all electrodes were included in all subsequent analysis steps following re-referencing. Bad electrodes were determined by voltage thresholding.

Custom Python codes were used to process the EEG data. We applied a high pass filter at 0.1 Hz, followed by a notch filter at 60 Hz and harmonics of 60 Hz to each participant’s raw EEG data. We then performed three preprocessing steps^51^ to identify electrodes with severe artifacts. First, we calculated the mean correlation between each electrode and all other electrodes as electrodes should be moderately correlated with other electrodes due to volume conduction. We z-scored these means across electrodes and rejected electrodes with z-scores less than -3. Second, we calculated the variance for each electrode as electrodes with very high or low variance across a session are likely dominated by noise or have poor contact with the scalp. We then z-scored variance across electrodes and rejected electrodes with a *|*z*| ≥* 3. Finally, we expect many electrical signals to be autocorrelated, but signals generated by the brain versus noise are likely to have different forms of autocorrelation. Therefore, we calculated the Hurst exponent, a measure of long-range autocorrelation, for each electrode and rejected electrodes with a *|*z*| ≥* 3. Electrodes marked as bad by this procedure were excluded from the average re-reference.

We then calculated the average voltage across all remaining electrodes at each time sample and rereferenced the data by subtracting the average voltage from the filtered EEG data. The average number of bad electrodes identified for each participant was (M=2.41, SD=1.02). Although bad electrodes were excluded from the average re-reference calculation, all electrodes were included for every participant for all subsequent analyses. We used wavelet enhanced independent component analysis^52^ to remove artifacts from eyeblinks and saccades.

### EEG Data Analysis

We applied the Morlet wavelet transform (wave number 6) to the entire EEG time series across electrodes, for each of 46 logarithmically spaced frequencies (2-100 Hz;^28^). We analyzed both stimulus-locked and response-locked signals. After log-transforming the power, we downsampled the data by taking a moving average across 100 ms time intervals 2500 ms preceding to 2000 ms following either probe presentation or response. We slid the window every 25 ms, resulting in 177 time intervals (45 non-overlapping). Power values were then z-transformed by subtracting the mean and dividing by the standard deviation power. Mean and standard deviation power were calculated across all decision phase trials and across time points for each frequency.

### Pattern Classification Analyses

Pattern classification analyses were performed using penalized (L2) logistic regression, implemented via the sklearn linear model module in Python and custom python code^32^. For all classification analyses, classifier features were composed of spectral power across 63 electrodes and 46 frequencies. Before pattern classification analyses were performed, we completed an additional round of z-scoring across features (electrodes and frequencies) to eliminate trial-level differences in spectral power^28^. Thus, mean univariate activity was matched for all conditions and trial types. We extracted “classification accuracy” which indicates the ability of the classifier to successfully guess condition labels (e.g. task label, “episodic,” “semantic,” and “perceptual”). We generated data-derived chance accuracy values for statistical evaluation of classifier performance by shuffling the condition labels of interest (e.g. “episodic,” “semantic,” and “perceptual”) and then calculating classification accuracy based on these shuffled labels. We repeated this procedure 1000 times for each participant and then averaged the 1000 shuffled accuracy values for each participant. These mean values were used as participant-specific empirically derived measures of chance accuracy. We also extracted “classifier evidence,” a continuous value reflecting the logit-transformed probability that the classifier assigns a particular label (e.g. encode, retrieve) to each trial. Classifier evidence served as a trial-specific, continuous measure of mnemonic state, task, or feature information.

### Cross-study Mnemonic State Analyses

We developed and validated a cross-participant mnemonic state classifier based on already collected data from participants (N=143,^19, 27^) who completed a mnemonic state task. In this paired-objects task, participants view two lists of images of common objects. All List 1 objects are new. All List 2 objects are also new, but are categorically associated to a List 1 object. For example, if List 1 contains an image of a bench, List 2 would contain an image of a different bench. During List 1, participants are instructed to encode each object. During List 2, however, each trial contains an instruction to either encode the current object (e.g., the new bench) or to retrieve the corresponding object from List 1 (e.g., the old bench). We performed within-participant leave-one-run-out classification (penalty=1) on patterns of spectral power across 63 electrodes and 46 frequencies to identify the participants who show robust classification accuracy. We used data-derived chance classification accuracy values to identify a subset of these participants (N=57) for whom true classification accuracy was at or above the 90th percentile of permuted chance values. Next, we conducted cross-validated classification (penalty parameter=0.0001) on the selected participants to validate our mnemonic state classifier. We obtained classification accuracy of 59.29% which is significantly above chance (*t* _56_=7.667, *p<*0.0001), indicating that the cross-participant mnemonic state classifier can distinguish memory encoding and memory retrieval states. We applied the cross-participant mnemonic state classifier to the stimulus- and response-locked decision phase data. Due to the structure of our classifier, positive classifier evidence values indicate greater evidence for the retrieval state and negative classifier evidence values indicate greater evidence for the encoding state. This approach reflects our assumption – based on theoretical models of memory brain states^17, 53^ – that encoding and retrieval exist along a continuum.

### Task Decoding Analyses

We conducted two cross-validated cross-participant pattern classification analyses on spectral power averaged across either the 2000 ms delay interval or the 500 ms preceding the response during the probe interval. We used leave-one-participant-out cross-validated decoding to avoid potential contributions of run number to decoding accuracy. We extracted classification accuracy which represented whether the classifier successfully guessed the correct task label (episodic, semantic, perceptual) for a given trial. We conducted two cross-interval classification analyses in which the classifier was trained on data from one interval (delay or probe) and tested on data from the other interval.

### Feature Decoding Analyses

We conducted six cross-validated within-participant pattern classification analyses on spectral power averaged across the 500 ms preceding the response during the probe interval. The six classifiers were trained to decode one set of features (e.g. old vs. new) that were either task-relevant (e.g. data from the episodic block) or task-irrelevant (e.g. data from both the perceptual and semantic blocks). We randomly subsampled 75% of the decision phase data for training and held out the remaining 25% for testing with equal numbers of trials (64 for task-relevant, 128 for task-irrelevant) from the episodic and perceptual feature conditions (old, new; uppercase, lowercase). We performed additional subsampling to balance the number of small vs. big items for the semantic feature decoders. We repeated the subsampling procedure for 100 iterations. We calculated the average classification accuracy values across iterations for each participant.

We conducted within-participant pattern classification analyses on probe-locked spectral power for each of 100 ms time intervals from probe onset (0 ms) to 1000 ms following probe onset. We used probe-locked data as we were interested in the earliest time point relative to stimulus onset when classifier evidence maximally dissociates the two feature conditions. We selected 1000 ms as the final time point based on the finding that the average median reaction time on correct trials across all tasks was 837.0 ms (SD=143.0 ms). The probe-locked classifiers were structured in the same manner as the response-locked classifiers with subsampling to equate trials per condition. Following pattern classification, we extracted the peak time interval for each classifier for each participant. Specifically, we obtained classification accuracy across all trials within a condition (e.g. old/new decoding accuracy for episodic task trials) for each time interval. The peak decoding time interval is that for which decoding accuracy was maximal.

### Statistical Analyses

We used repeated-measures ANOVAs (rmANOVAs) and paired-sample *t* -tests to assess the effect of task (episodic, semantic, perceptual) on response accuracy and reaction time. We used rmANOVAs to assess the impact of task and time interval on mnemonic state evidence.

We used paired-sample *t* -tests to compare classification accuracy across participants to chance-level decoding accuracy, as determined by permutation procedures. For the feature decoders, for each participant, we shuffled the condition labels of interest (e.g., old and new) and then calculated classification accuracy. We repeated this procedure 1000 times for each participant and then averaged the 1000 shuffled accuracy values for each participant. These mean values were used as participant-specific empirically derived measures of chance accuracy. We utilized the same general procedure to assess leave-one-participant-out task classification accuracy.

We used Wilcoxon-rank sum tests to compare the median peak decoding time intervals across task and task-relevance conditions^54^.

We used Pearson correlations to link the retrieval state with task and feature information. We conducted within-participant trial-level correlations between retrieval state evidence and task or feature evidence, separately for each task or feature. We Fisher-Z transformed the resulting rho values and averaged the z-Rho values across tasks or features. We used one-sample *t* -tests to assess the extent to which trial level mnemonic state evidence and task or feature evidence are correlated.

We report effect sizes as partial eta squared (*η_p_*^2^), Cohen’s d, and rank-biserial correlation coefficient. We conducted Bayes Factor analysis using the Bayes Factor package (version 0.9.12-4.7) in R (version 4.5.1) with the default prior settings and specifically the linear model function lmBF to compare models.

## Funding Statement

This work was supported by a grant from the National Institutes of Health (NINDS R01 NS132872, PI: N.M.L.).

## Data availability

The datasets generated in the current study will be made available on the Open Science Foundation (OSF) via the Long Term Memory Lab website (http://longtermmemorylab.com/publications/) upon publication.

## Code availability

All experimental codes used for data collection and data analysis will be made available on the Open Science Foundation (OSF) via the Long Term Memory Lab website (http://longtermmemorylab.com/publications/) upon publication.

## Notes

### Competing Interest Statement

The authors have declared no competing interest.

